# Neural decoding of gait phase information during motor imagery and improvement of the decoding accuracy by concurrent action observation

**DOI:** 10.1101/2020.08.19.258210

**Authors:** Hikaru Yokoyama, Naotsugu Kaneko, Katsumi Watanabe, Kimitaka Nakazawa

## Abstract

Brain decoding of motor imagery (MI) is crucial for the control of neuroprosthesis, and it provides insights into the underlying neural mechanisms. Walking consists of stance and swing phases, which are associated with different biomechanical and neural control features. However, previous studies on the decoding of the MI of walking focused on the classification of more simple information (e.g., walk and rest). Here, we investigated the feasibility of electroencephalogram (EEG) decoding of the two gait phases during the MI of walking and whether the combined use of MI and action observation (AO) would improve decoding accuracy. We demonstrated that the stance and swing phases could be decoded from EEGs during AO or MI alone. Additionally, the combined use of MI and AO improved decoding accuracy. The decoding models indicated that the improved decoding accuracy following the combined use of MI and AO was facilitated by the additional information resulting from the concurrent cortical activations by multiple regions associated with MI and AO. This study is the first to show that decoding the stance versus swing phases during MI is feasible. The current findings provide fundamental knowledge for neuroprosthetic design and gait rehabilitation, and they expand our understanding of the neural activity underlying AO, MI, and AO+MI of walking.

**Significance Statement:** Brain decoding of detailed gait-related information during motor imagery (MI) is important for brain-computer interfaces (BCIs) for gait rehabilitation. However, previous knowledge on decoding the motor imagery of gait is limited to simple information (e.g., the classification of “walking” and “rest”). Here, we demonstrated the feasibility of EEG decoding of the two gait phases during MI. We also demonstrated that the combined use of MI and action observation (AO) improves decoding accuracy, which is facilitated by the concurrent and synergistic involvement of the cortical activations by multiple regions for MI and AO. These findings extend the current understanding of neural activity and the combined effects of AO and MI and provide a basis for developing effective techniques for walking rehabilitation.

## Introduction

Walking is one of the most common and essential daily human movements. Thus, gait rehabilitation after neurological disorders has been extensively investigated (1, 2). Recent studies have demonstrated that brain-computer interface (BCI)-based gait neurorehabilitation systems do not only support patient movements; they can also induce neurological functional recovery (3, 4). Neural decoding techniques, which predict the motor state of a human from brain signals, are a crucial part of BCI systems (5). In addition to the essential role in BCI, neural decoding can provide insights into the underlying neural mechanisms of target movements (6).

Previous gait rehabilitation studies on BCI were based on neural decoding of simple motor imagery (MI) information (e.g., discrimination of brain states between “walk” and “rest” (3, 7) and between “left leg” and “right leg” (4)). However, studies on upper-limb movements have succeeded in decoding more detailed and functional information of MI, such as hand postures (8, 9) and movement directions (10). Based on the upper-limb studies, more functionally related motor information may be decoded from brain activity during MI of walking.

Walking consists of two main phases, swing and stance phases, which are associated with different muscular functions (11). It has been reported that gait phase-dependent patterns of the functional electrical stimulation (FES) of the leg muscles improve walking performance in individuals with neurological disorders (12, 13). In addition to the functional support, FES timed with voluntary effort effectively induces neurological recovery (14). Therefore, neural decoding of gait phases may contribute to the development of novel gait rehabilitation based on BCI-FES systems, which will effectively evoke neural plasticity.

Although MI and action observation (AO) have traditionally been recognized as two distinct rehabilitation techniques, their combined use (MI+AO) can be more effective in rehabilitation; it can induce increased cortical activity compared with AO or MI alone (15, 16). MI mainly activates the primary motor cortex (M1) and supplementary motor area (SMA), which are considered to be involved in mental motor simulation (17, 18). On the other hand, AO is associated with the activation of the premotor, inferior frontal gyrus (IFG), temporal, posterior parietal, and occipital cortices, which may be involved in visual processing and the mirror neural system for action understanding (19, 20). During hand movements, it has been reported that MI+AO induces stronger activations than MI or AO alone in cortical regions related to mirror neuron and motor systems (21, 22). Regarding walking, a recent transcranial magnetic stimulation study demonstrated enhanced corticospinal excitability during MI+AO compared with AO alone (23). Given the combined effect of MI and AO, MI+AO may contribute to higher decoding accuracy of gait phases than MI or AO alone.

Therefore, we here tested the feasibility of the neural decoding of gait phases during MI and/or AO and investigated the combined effect of MI and AO. We demonstrate that gait phases can be decoded from cortical signals, and the decoding accuracy is higher during AO+MI than MI or AO alone. We also show that multiple cortical regions, including motor, visual, and mirror neuron systems, contribute to the higher decoding accuracy during AO+MI. This study demonstrates the feasibility of decoding the stance and swing phases during MI and the effectiveness of the combined use of MI and AO for decoding. Our results also provide the fundamentals for understanding the neural mechanisms underlying the MI and AO of walking and developing an effective BCI system for gait rehabilitation.

## Results

### Overview of the study

Thirteen healthy males participated in this study. They performed the following three tasks: MI, AO, and AO+MI (Fig. 1B–D). For AO+MI (Fig. 1A), they were instructed to observe the walker’s right leg and imagine that they were walking like the person. For AO (Fig. 1B), the participants were instructed to observe the walker’s right leg and not imagine anything for one minute. For MI (Fig. 1C), the participants were instructed to imagine that they were walking following the instructions for gait events (stance and swing phases) presented on an LCD monitor. EEG signals were recorded from 62 channels, and artifacts in the EEGs were removed by procedures based on independent component analysis (ICA). EEG data from 46 channels located in the frontal area (Figure 1D) were used for subsequent analysis because it was suggested that eye movement artifact may persist in the frontal electrodes even after the ICA method in free viewing experiments (24). Using the time-series information of the gait phase and EEG signals, we designed neural decoders to classify cortical activity into the stance or swing phases. See Figure 2 for an overview of our decoding methodology.

**Figure 1.**
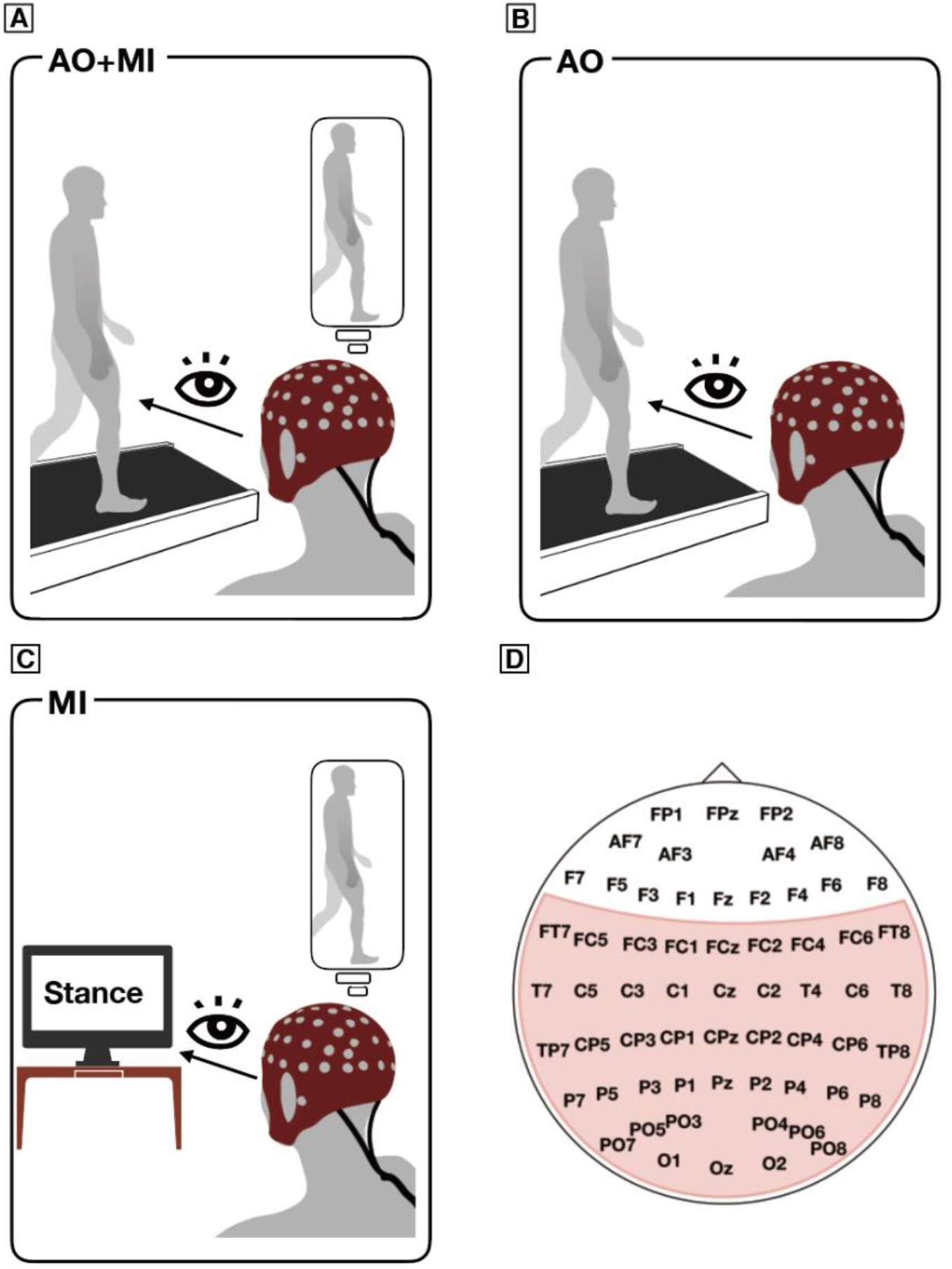
Experimental setups (A–C) and EEG electrode layout (D). (A–C) Experimental setups for the following three conditions: action observation of walking with motor imagery (AO+MI, panel (A)), action observation of walking (AO, panel (B)), and motor imagery (MI, panel (C)). (D) Sixty-two channels of electrodes based on the international 10-10 system, which were used for the EEG preprocessing, are shown. Given the effects of eye movements on the frontal electrodes, electrodes in the red area were used for decoding analyses.

**Figure 2.**
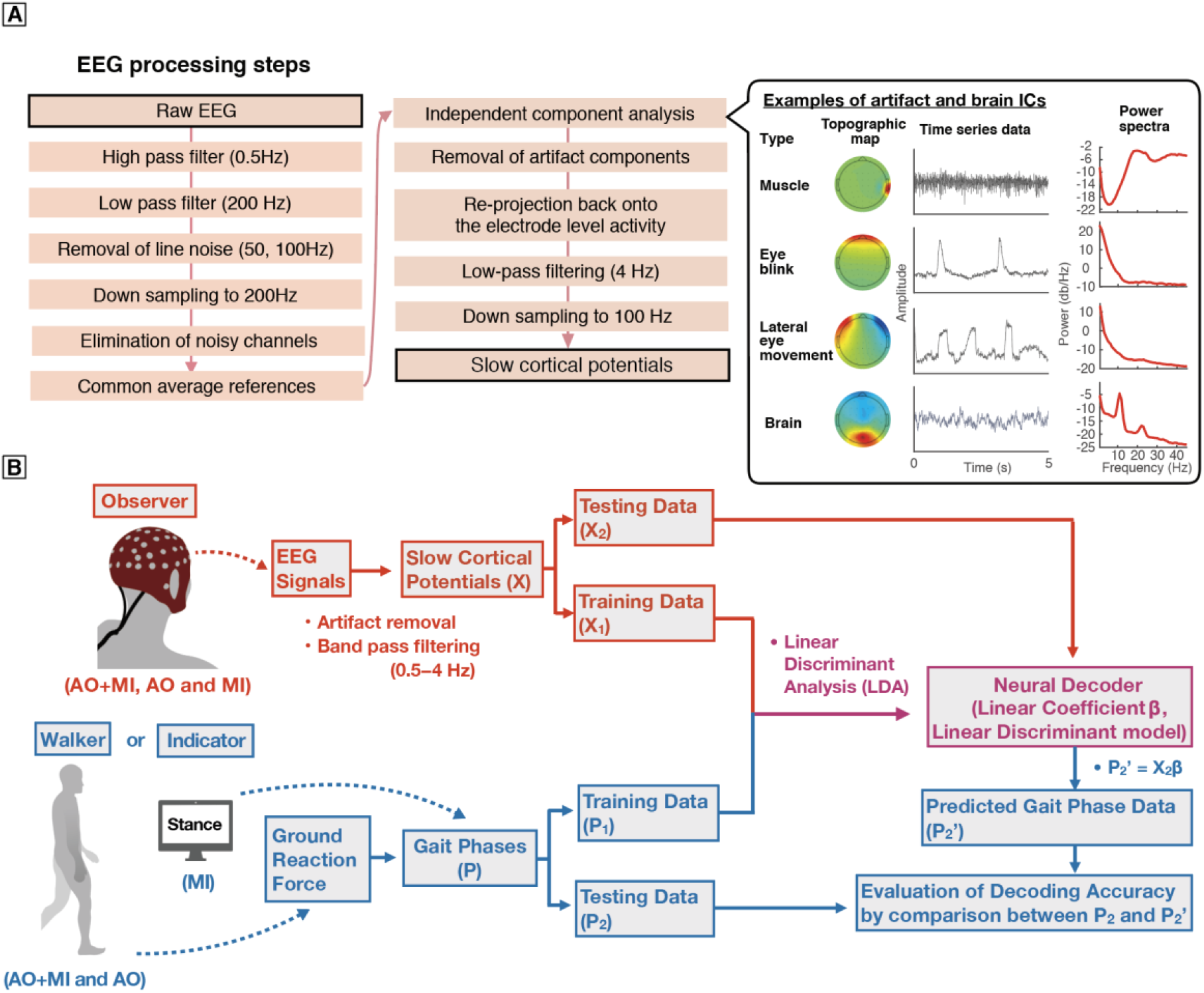
Schematic flow chart illustrating EEG processing steps (A) and neural decoding of gait phases (B). (A) Flowchart of EEG processing steps showing filter characteristics, down-sampling rate, and procedures for artifact removal based on the independent component analysis. In the right part, typical examples of topographic maps, time-series signals, and power spectra for artifacts and brain components are shown. (B) Flowchart of neural decoding of gait phases. Gait phases of a walker or an indicator and slow cortical potentials obtained from EEG signals were trained using linear discriminant analysis (LDA). The neural decoder (linear coefficients) obtained from the training data were used to predict the gait phases of the testing data.

### Decoding accuracy of the neural decoders

In preparation for neural decoding, EEG signals were band-pass filtered in the delta band (0.5-4Hz). The filtered signals, which are called slow cortical potentials, were confirmed to be informative for decoding actual movements (25–29) and motor intention (30–32). We used regularized linear discriminant analysis (rLDA) to decode the gait phases from cortical signals, as used in previous EEG decoding studies (33–35). Figure 3 shows typical examples of actual and decoded gait phases during AO+MI (Fig. 3A), AO (Fig. 3B) and MI (Fig. 3C) by a participant. In this participant, the time course of classification from the neural decoders showed alternating patterns between the swing and stance phases roughly matching the actual gait-phase time-series data of all three conditions (AO+MI, AO, MI). The decoding accuracies of the neural decoders were in the following order: AO+MI > AO > MI.

**Figure 3.**
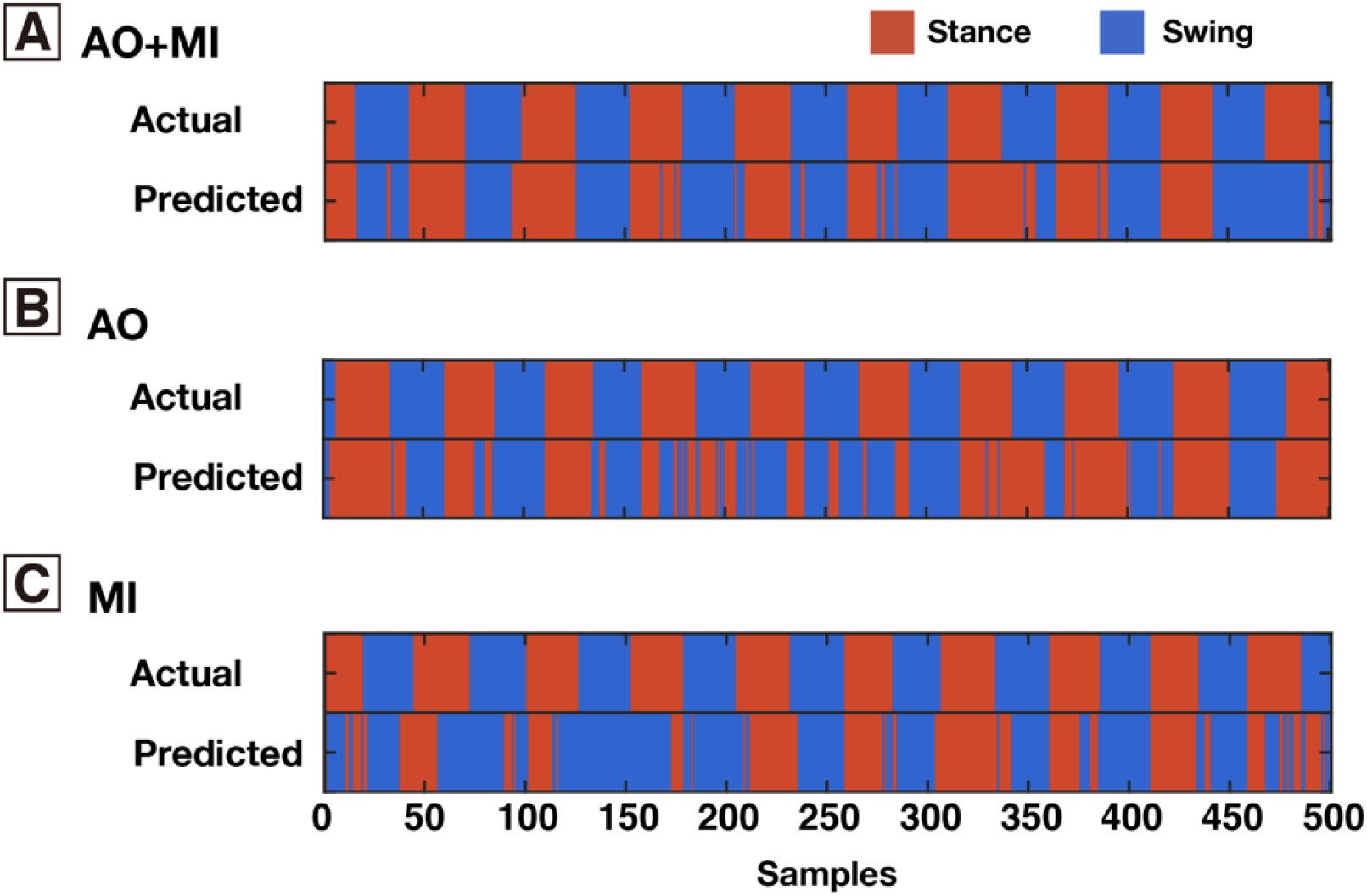
Typical examples of actual and decoded gait phases of a participant. Examples of 500 consecutive samples of the decoded results in each task are shown. Red and blue areas indicate the stance and swing phases, respectively. The results during AO+MI (A), AO (B), and MI (C) tasks are shown.

To test if the decoding accuracies of the neural decoders exceeded the chance level, we examined the confidence intervals using a randomization test (36). The labels of the feature vectors (stance or swing phases) were randomly permuted over the time points for each subject and condition. We generated 500 surrogate datasets and evaluated the upper end of 95% of the distribution of their decoding accuracies (mean chance levels shown in Fig.4). The decoding accuracies of the original EEGs exceeded the chance level of the surrogate datasets for all the conditions and the participants (see Table S1 for all individual values).

Figure 4 shows the differences between the decoding accuracies of the three conditions (Fig. 4). The decoding accuracy of AO+MI (72.2±7.5 %: mean±standard deviation [SD]) was significantly higher than those of AO (66.3±8.7 %) and MI (62.3±7.0 %) (ANOVA: F2,24 = 8.17, p = 0.0020; paired t-test with FDR correction: p = 0.011 for AO+MI vs. AO, p = 0.0016 for AO+MI vs. MI). There was no significant difference between the decoding accuracies of AO and MI (paired t-test with FDR correction, p = 0.26). The detailed statistical values are summarized in Tables S2.

**Figure 4.**
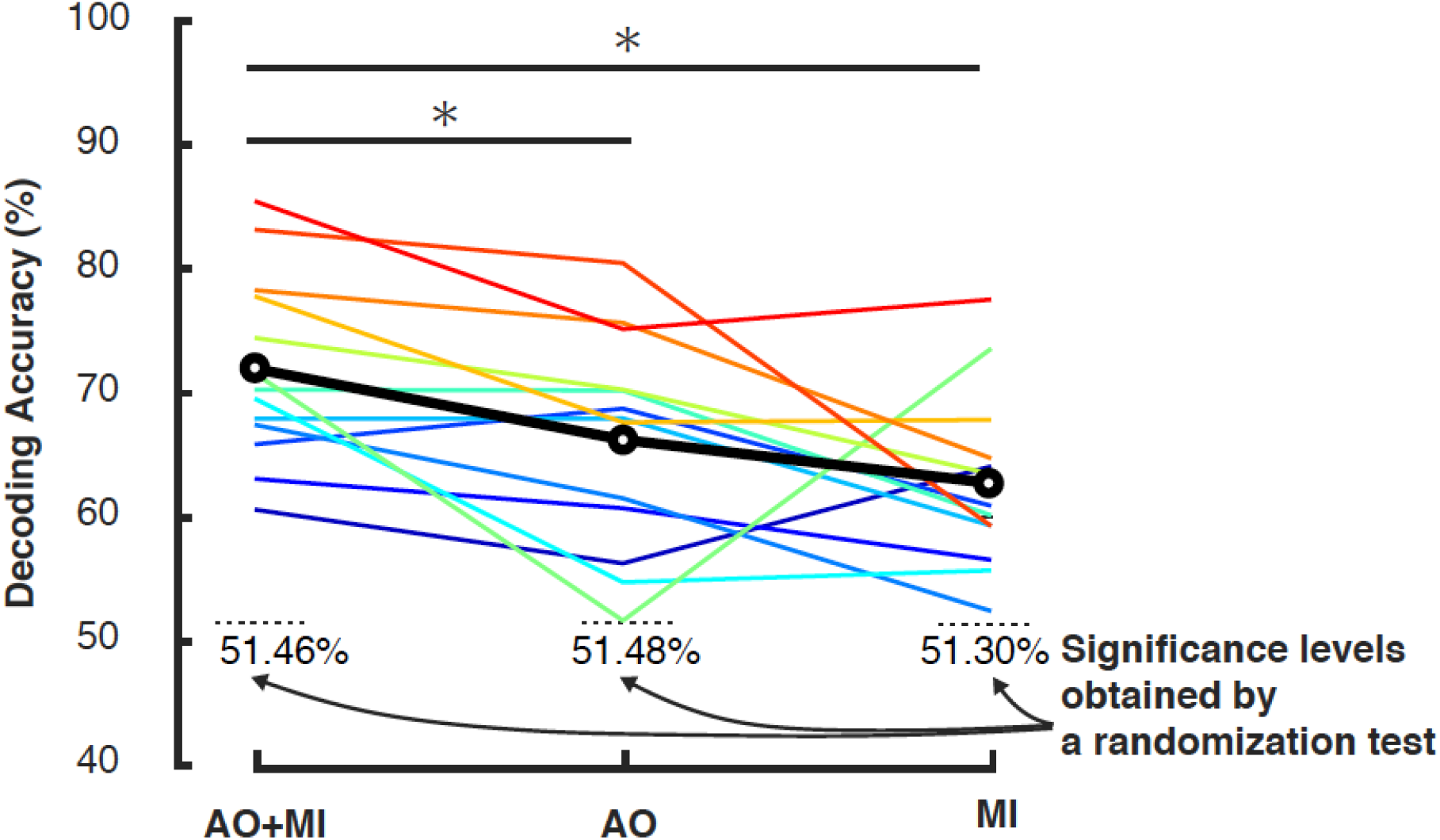
Decoding accuracies for three tasks (AO+MI, AO, and MI). Black plot indicates the mean values across participants. Colored plots indicate individual data. Horizontal black dash lines indicate participant means of significance (chance) level for each condition obtained by a randomization test. All the individual data exceeded the chance level values in all the tasks. Asterisks indicate significant differences (* p < 0.05, paired t-test with FDR correction, See also Tables S1 and S2 for detailed statistical values).

We calculated the decoding accuracy of each label (swing phase and stance phase) independently and summarized the results in a confusion matrix (Fig. 5). The decoding accuracy was significantly higher during the stance phase than during the swing phase for all three conditions (p < 0.001, paired t-test, see Tables S3 for detailed statistical values).

**Figure 5.**
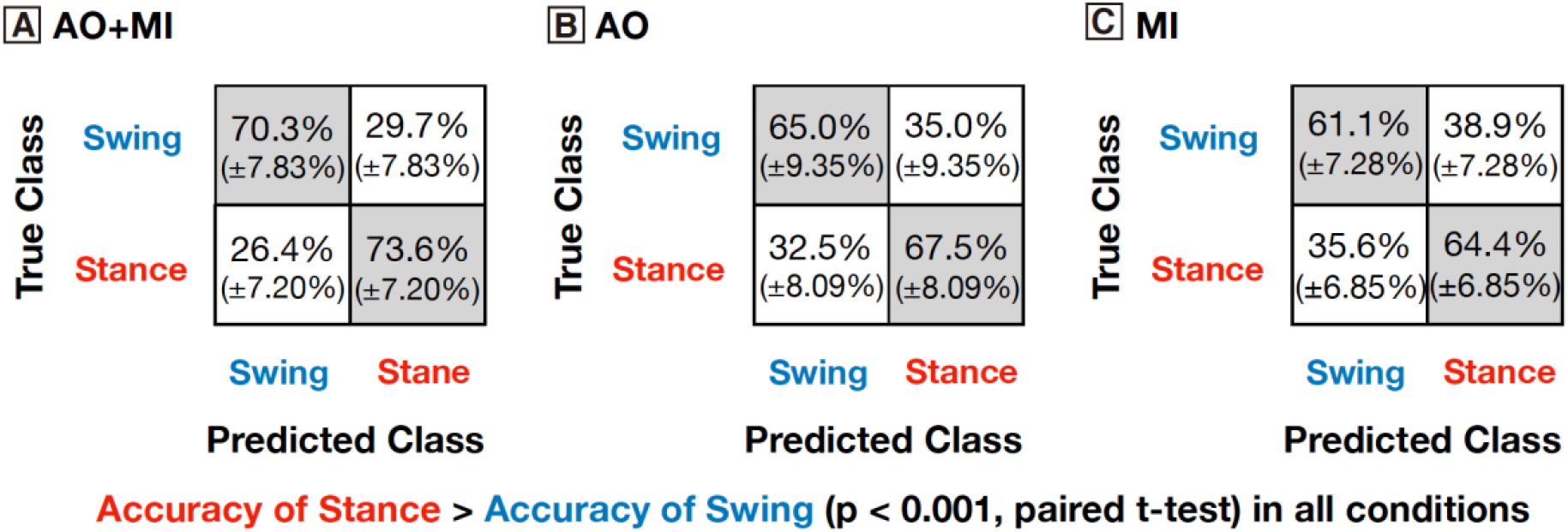
Confusion matrices (in %) of the swing and stance phases. Gourp mean accuracies and the standard deviations are shown for all three tasks. The decoding accuracy was significantly higher during the stance phase than during the swing phase in all three tasks (p < 0.05, paired t-test, See also Tables S3 for detailed statistical values).

### Contributions of Electrodes to Neural Decoding

To evaluate the topographic contributions of cortical activities to the decoding of the gait phases, we calculated the percentage contribution of each electrode from the weights of the decoding model based on previous studies (29, 35, 37). Figure 6 shows the mean data and all individual topographic contributions of weights from the neural decoding models. For AO+MI, the left frontotemporal and occipital areas showed higher contributions based on the visual inspection of the mean topographic map of the participants (Fig. 6A). Higher contributions by these cortical regions were demonstrated by most of the individual data (Fig. 6B). In the group mean topographic map for AO+MI, the central midline area also showed a high contribution, but it was slightly lower than those of the left frontotemporal and parietal regions (Fig. 6A). In this cortical area, some participants (e.g., ID 1, 2, 4, and 9) showed high contributions, while others did not (e.g., ID 8 and 10) (Fig. 6B). For AO, the group mean topographic patterns showed higher contributions by the left frontotemporal and occipital areas, as demonstrated for AO+MI (Fig. 6C). Most of the participants showed high contributions by these areas (Fig. 6D). For MI, higher contributions of the central midline and occipital areas were observed in the group mean topographic patterns (Fig. 6E). For the individual data, most participants showed a high contribution of the occipital area (Fig. 6F). Some participants (e.g., ID 2, 12, and 13) showed a high contribution of the central midline area (Fig. 6F).

**Figure 6.**
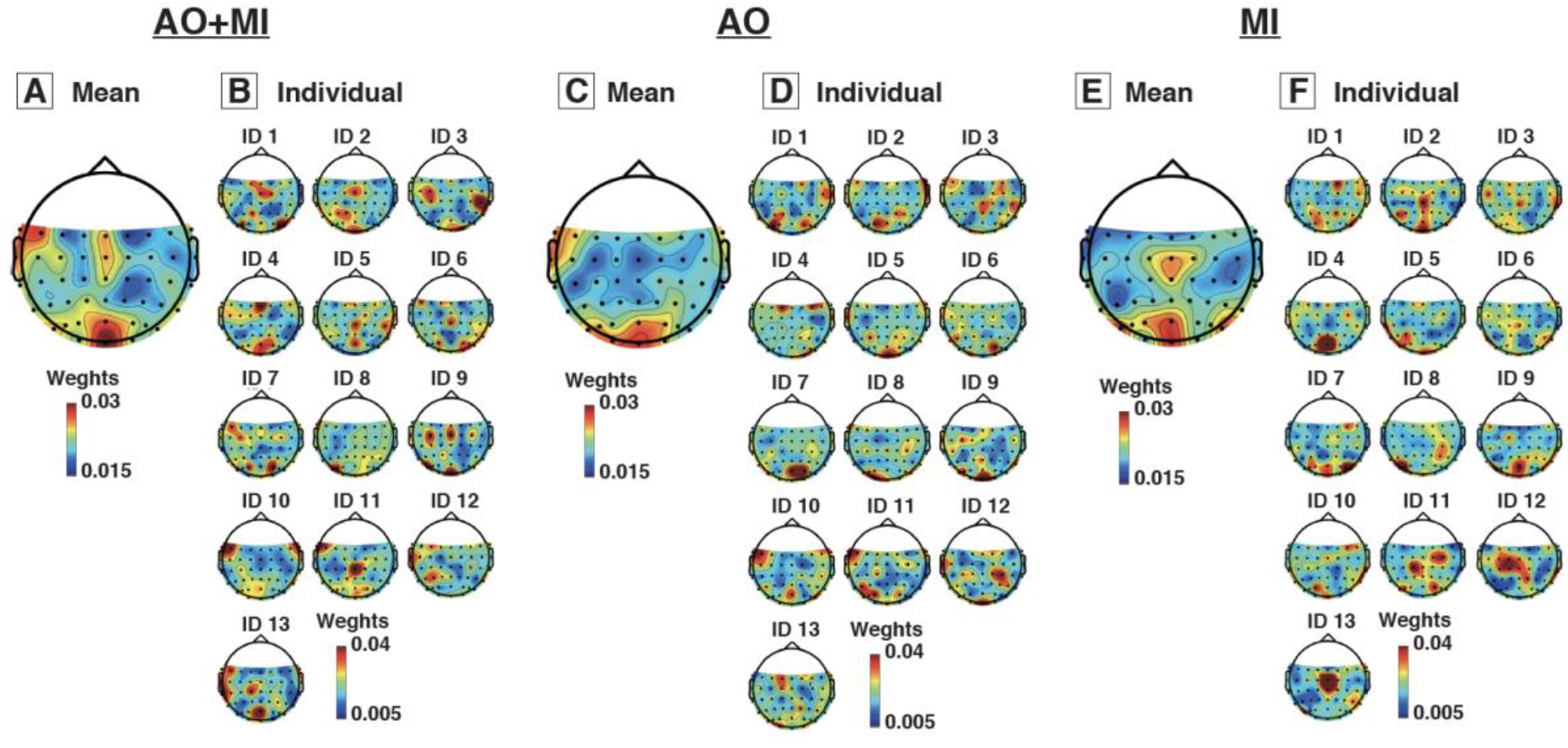
Topographic distribution for weights of neural decoding models. Group means and individual data are separately shown for the AO+MI (A), AO (B), and MI (C) tasks.

The above-mentioned results suggested that different cortical regions were involved in decoding during each condition. To quantitatively examine the contributions of each cortical region during each condition, we divided the electrodes into 11 regions of interest (ROIs) (Fig. 7A). Subsequently, the average contributions within each ROI were compared to the mean contribution across all the electrodes as a baseline (0.217 = 1/46 electrodes) (Fig. 7B−7D). For the AO+MI condition, the mean weights of the left anterotemporal, left occipital, and right occipital ROIs were significantly greater than the baseline weight (p < 0.05, permutation test with FDR correction) (Fig. 7B). For the AO condition, as in AO+MI, the mean weights of the left anterior-temporal and left and right occipital ROIs were significantly greater than the baseline weight (p < 0.05, permutation test with FDR correction) (Fig. 7C). On the other hand, the mean weights of the left and right frontocentral and left parietal ROIs were significantly lower than the baseline weight (p < 0.05, permutation test with FDR correction) (Fig. 7C). For the MI condition, the left and right occipital ROIs showed significantly larger weights than the baseline, while the left anterotemporal ROI demonstrated significantly lower weights (p < 0.05, permutation test with FDR correction) (Fig. 7D). In summary, the decoding models of AO+MI were mainly based on information from the cortical regions, which had higher contributions in the models for MI and AO.

**Figure 7.**
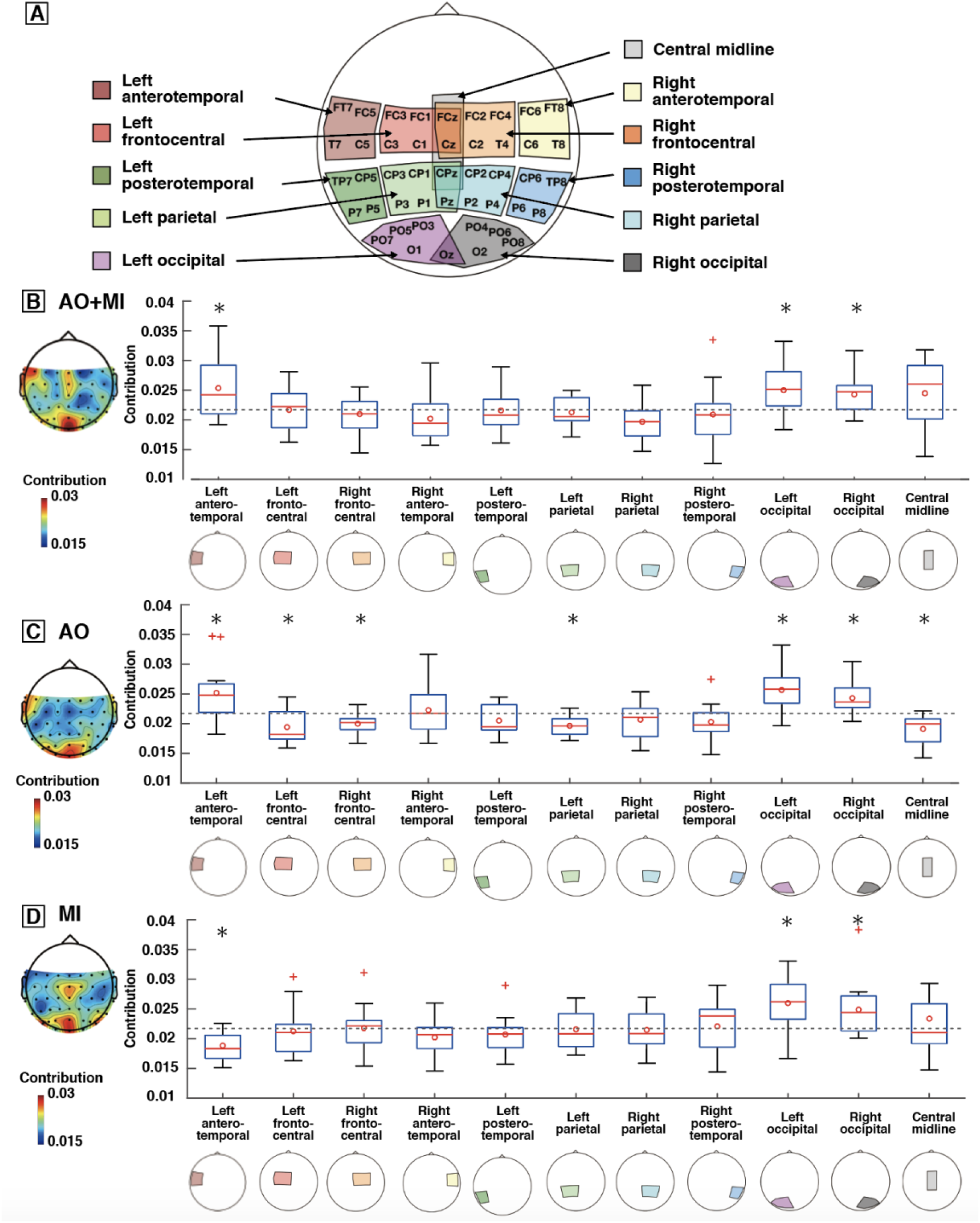
Weights of decoding models in each region of interest (ROI). (A) Scalp map indicating the electrodes included in each ROI for investigating the contributions of the cortical regions to the decoding of the gait phases. (B−D) Boxplots showing the weights of each ROI during AO+MI (panel B), AO (panel C), and MI (panel D). The edges of the box indicate the 75th percentiles. Red lines and circles indicate the median and mean, respectively. Whiskers mark the range of values. Outliers are plotted as crosses. The mean weight of all the electrodes is shown as the horizontal dash lines. The asterisks indicate significant differences among the mean weights of all the electrodes (p < 0.05, permutation test with FDR correction. See also Tables S4–S6 for detailed statistical values). The mean topographic distribution for weights across participants (presented in Fig. 6) is shown on the left in each panel.

## Discussion

Our results demonstrated that gait phases could be significantly decoded from cortical slow waves during MI+AO, MI, and AO, and the decoding accuracy during MI+AO was greater than that of MI or AO alone (Fig. 4). In addition, the higher decoding accuracy during AO+MI was facilitated by multiple cortical regions including the motor, visual, and left frontotemporal regions based on the decoding models (Figs. 6 and 7).

### Higher effectiveness of the combined use of AO and MI than AO or MI alone

The decoding accuracy of the gait phases was higher during MI+AO than during MI or AO alone (Fig. 4). So far, the effectiveness of the combined use of AO and MI for motor learning and rehabilitation has been studied. In motor learning research, more improvements of the AO+MI intervention than AO or MI alone have been reported for golf putting (38), dart-throwing (39), and the bicep curl strength test (40). Recently, it was also reported that AO+MI can improve the motor performance of individuals after stroke (41) and total hip arthroplasty (42), although the rehabilitation outcome of AO+MI was not compared to that of AO or MI.

In the field of neural engineering, brain activities during AO and MI have been used as information sources for neural decoding of movements for developing BCI systems for individuals with paralysis (43, 44). Regarding the effectiveness of the combined use of AO and MI for neural decoding, a recent study using fMRI demonstrated that the accuracy of decoding for classifying brain states among different upper-limb motor tasks was higher during AO+MI than MI alone (45). However, the following points were still open for question: (1) improvement of decoding accuracy by the combined use of AO and MI compared to AO alone and (2) combined effects of MI and AO on neural decoding of gait-related information. Thus, the higher decoding accuracy of gait phases during MI+AO than MI or AO alone provide further evidence for the effectiveness of AO+MI for neural decoding of gait information in addition to other evidence showing the effectiveness of AO+MI for motor learning and rehabilitation as well as the neural decoding of the upper limb movements.

### Topographic cortical contribution to the decoding of the gait phase

We examined the topographic contributions of cortical activity to decoding based on the weights of the decoding models (Fig. 7). We found higher contributions of the left and right occipital ROIs during AO (Fig. 7C). The occipital region activity during AO would be involved in visual processing (15). The large contribution from the occipital region during AO in this study is consistent with findings from previous fMRI studies, which demonstrated high activity of the occipital regions during AO of hand movements (46) and walking (47). Also, the left anterotemporal ROI showed higher contributions to decoding (Fig. 7C). The anterotemporal ROI spatially overlaps with a part of the mirror neuron system, including the IFG and ventral premotor cortex (48, 49). The mirror neuron system is thought to be involved in AO for the understanding of the observed movements (15, 20, 50). In the present study, only the left side of the anterotemporal ROI showed a significantly higher contribution (Fig. 7C). Regarding the motor cognition of the left hemisphere, apraxia, a disorder that presents with deficits in the performance of purposeful movements such as imitation, is frequently observed in patients with left hemisphere damage (51). Also, it has been demonstrated that limb-apraxic patients had an impairment in gesture comprehension and the severity of the comprehension was correlated with the size of the lesion of the left IFG (52). Thus, the left hemisphere may be important for understanding actions. The lateralized functions are possibly related to the high contribution of the left anterotemporal ROI during AO in the present study. On the other hand, ROIs in the sensorimotor area (left and right front-central, left parietal and central midline ROIs) showed low contributions (Fig. 7C). This indicates that cortical activity in the sensorimotor area does not contain much gait phase information during the AO of walking.

Regarding cortical involvement during MI, two different processes are considered to be typically involved in MI; they are termed as the visual-motor and the kinesthetic motor imagery processes (53). The visual-motor imagery process consists of the visualization of the movements of the limbs, which does not require any special training or fine kinesthetic sensation; the kinesthetic motor imagery involves the feeling of limb movements through the kinesthetic sensation coming from the limbs, which is usually highly achieved by athletes (54). The high contribution of the occipital cortex (i.e., visual cortex) during MI in the present study (Fig. 7B) was probably related to the visual-motor imagery process. Some previous studies demonstrated the involvement of the occipital cortex in the visual-motor imagery process (55, 56). As another possible reason, the high contribution of the occipital cortex may have been related to the different visual cues displayed by the monitor for the stance and swing phases. The contribution of the sensorimotor area ROIs was low during AO as mentioned above (Fig. 7C), but it was not during MI (Fig. 7B). The difference may have been related to the involvement of the sensorimotor regions in the kinesthetic motor imagery process during MI (57). Also, the insignificantly high contribution of the sensorimotor area ROIs (Fig. 7B) may be attributed to the large inter-participant variability of kinesthetic imagery ability (54, 55). In line with this possibility, some participants showed a high contribution of the sensorimotor areas; the others demonstrated low contributions (Fig. 6B).

For AO+MI, we observed higher contributions of the left and right occipital and the left anterotemporal ROIs (Fig. 7A). As with MI, the contribution of the sensorimotor regions, which was low during AO, was not significantly different from the mean contribution across the electrodes during AO+MI. The difference between the contribution patterns during the tasks suggests that the underlying cortical processes were different for AO and MI, and these processes were concurrently involved during AO+MI. This may explain the higher decoding accuracy during AO+MI than AO or MI alone (Fig. 4). Also, a recent study suggested that the MI-component of AO+MI works as the motor simulation of a participant’s movements, whereas the AO-component of AO+MI works as an external visual scaffold for MI (58). Therefore, visual guiding may improve the mental motor simulation during AO+MI. This synergistic effect may also have contributed to the higher decoding accuracy during AO+MI (Fig. 4).

### Difference between the decoding accuracies of the stance and swing phases

The decoding accuracy was higher during the stance phase than during the swing phase of all tasks (Fig. 5 and Table S3). So far, the swing phase has been thought to be critical for cortical involvement in muscle control during walking (59, 60). However, recent studies demonstrated significant corticomuscular connectivity during the stance phase (61, 62). Additionally, sensory inputs from the hip flexors and ankle extensors during the stance phase are essential for determining the timing of the gait-phase change (63, 64). Load-related sensory information from the sole is provided only during the stance phase. The stance phase-specific sensory information probably enhances motor simulation and action understanding. Thus, cortical involvement in muscle control and sensory processing for the stance phase may be associated with higher decoding accuracy during the stance phase.

### Possible applicability to Brain-Computer Interfaces

Previous studies on BCI have been based on neural decoding of simple MI information (e.g., the classification between “walk” and “rest” (3, 7) and between “left leg” and “right leg” (4)). On the other hand, our results suggest that it is feasible to decode more functionally related information, which is the gait-phase information. The neural decoding of gait phases will contribute to developing BCI-FES systems because timing information of the gait phases is vital for FES for gait rehabilitation (12, 13). Further, FES timed with voluntary effort, which can be achieved by the BCI-FES systems, can effectively induce neurological recovery (14). Thus, our results may contribute to the future development of BCI systems for improving gait performance and inducing neural recovery.

### Limitations of the Study

Although we discussed the applicability of our results to BCI systems, there are several issues with applying our findings. The mean decoding accuracy (slightly above 70%, Fig. 4) is not enough for effective BCI-FES systems. Although we used LDA, considering the high interpretability of the neural decoding models, non-linear models will improve the decoding accuracy (65). Also, real-time decoding is necessary for BCI-FES systems. Additionally, we need to develop decoding models with considerations for cortical activity and EEG artifacts associated with participant movement and electrical stimulation.

### Conclusion

The results show that decoding the stance and swing phases during MI is feasible. We also showed that the combined use of MI and AO improves decoding accuracy. The improved decoding accuracy during AO+MI may have been derived from concurrent involvement of the cortical activations related to sensorimotor, visual, and action understanding systems associated with MI and AO. The results provide the fundamentals for developing effective BCI systems for gait rehabilitation and understanding the underlying neural activities during AO, MI, and AO+MI.

## Materials and Methods

### Participants

Thirteen healthy male volunteers (age, 22−32 years) participated in this study. Each participant provided written informed consent for participation in this study. The experiments were performed in accordance with the Declaration of Helsinki and with the approval of the ethics committee of the University of Tokyo.

### Experimental setup and design

We asked the participants to perform tasks for MI, AO, and AO+MI (Fig. 1B–D). Participants sat on a chair located approximately 1.5 m away from a treadmill (Bertec, Columbus, OH, USA) during AO and AO+MI tasks and a desk during the MI task. For the AO and AO+MI tasks, a healthy male walked on the treadmill at a fixed speed of 1.0 m/s. The person participated in all the experiments for all the participants. For the AO task, the participants were instructed to observe the walker’s right leg and not imagine anything for one minute. For the AO+MI task, they were instructed to observe the walker’s right leg and imagine that they were walking like the person for one minute. During the AO and AO+MI tasks, the participants observed the walker from a lateral view. For the MI task, the participants were instructed to imagine that they were walking following the instructions of gait events (stance and swing phases) presented by an LCD monitor on a desk. The monitor showed “stance” and “swing” alternately. The timing for switching between the two was determined in advance based on the average stance and swing duration of the participant during the AO and AO+MI tasks. For all the tasks, the participants were instructed to relax and concentrate during the recording. The duration was 1 minute for all the tasks. The participants performed the three different tasks six times in a random order (six minutes for each condition).

### Data collection

Three-dimensional ground reaction forces (GRF) were recorded from the walker with force plates beneath the right and left belts of the treadmill at a sampling rate of 1000 Hz. The GRF data were low-pass filtered (5 Hz cutoff, zero-lag Butterworth filter). MATLAB 2019a (MathWorks, Natick, MA, USA) was used to perform all the post-processing analyses offline.

EEG signals were recorded from 61 channels using an EEG cap (Waveguard original, ANT Neuro b.v., Enschede, Netherlands) according to the international 10-10 system layout (Fig. 1D) and an EEG amplifier (eego sports, ANT Neuro b.v., Enschede, Netherlands) at a sampling frequency of 500 Hz. Ground and reference electrodes were placed on AFz and CPz. Impedances of the electrodes were kept below 30 kΩ (10 kΩ in most electrodes), which was substantially lower than the recommended impedance (below 50 kΩ) for the high-impedance EEG amplifier.

### Data analysis

#### EEG pre-processing

In the current study, fluctuations in the amplitude of slow cortical potentials (0.5−4 Hz in the time domain) were used for the neural decoding analysis based on a similar methodology used in previous studies on actual movements (25–29) and motor intention (30–32). An overview of the EEG processing steps is shown in Figure 2.

We performed analyses of the EEG signals using custom-written programs in MATLAB 2019a (The MathWorks, Natick MA) with functions of EEGLAB 14.1b (66). An overview of the EEG pre-processing procedure is shown in Figure 2A. The EEG signals were high-pass filtered (cutoff frequency: 0.5 Hz, 4th order Butterworth filter) and low-pass filtered (cutoff frequency: 200 Hz, 4th order Butterworth filter). We reduced the 50 Hz line noise using the “cleanline” function in EEGLAB and resampled the signals to 200Hz. Subsequently, we checked the bad channels that had standard deviations higher than 1mV or a kurtosis above five standard deviations from the mean (29, 67). No EEG electrode was classified as a bad channel for all the participants. The EEG signals were re-referenced to a common average. The original reference electrode was recalculated as FCz, generating a total of 62 cortical electrodes.

Subsequently, each participant’s EEG signals were subjected to independent component analysis (ICA) to remove artifacts in the EEG signals (67, 68). From the extracted ICs, artifact ICs were visually identified and removed based on the following criteria: (1) muscle artifact: power in the 20–100 Hz was higher than that in the alpha bands (8–12 Hz) (69); (2) eye-blink and horizontal eye-movement artifact: power of the frequency below 3Hz was higher than that of the faster frequency – there was no clear power peak in the alpha (8–12 Hz) and beta (15–30 Hz) bands, and the topological map and the time-series data corresponded to previously reported features of these components (69, 70). Examples of the artifact and brain components are presented in Figure 2A. Only the remaining ICs were projected back onto the electrode-level activity. Regarding the eye movement artifact in free viewing experiments, a recent paper suggested that the ICA-based correction is effective in reducing eye activity artifacts, but the residual eye movement artifact may persist in the frontal electrodes (24). Since the AO+MI and AO were free viewing tasks in this study, we used 46 electrodes, except those located in the frontal area (Fig. 1D), in the following decoding analyses. Brain areas that are considered to be largely involved in motor imagery and observation (Motor, temporal, parietal, and visual cortical areas) were covered by the remaining electrodes (71, 72). The EEG signals were low-pass filtered with a 4th order Butterworth filter (cutoff frequency: 4 Hz) to obtain slow cortical potentials. The amplitude of each electrode was normalized to unit variance and zero mean. Finally, the EEG data were downsampled to 100Hz.

#### Neural decoding of gait phases

Figure 3 shows an overview of our decoding methodology. We designed neural decoders (linear discriminant model) that predict gait phases from slow cortical potentials. A time-embedded EEG dataset was constructed for each time point as a feature vector (i.e., predictors) in the linear discriminant analysis. The feature vector for each time point was constructed by concatenating the 21 lags (the current time point plus the 10 prior and 10 future [corresponding to 100 ms prior and 10 future]) for each channel into a single vector of length 21 × *N*_*electrode*_, where *N*_*electrode*_ is the number of EEG channels used for the linear discriminant analysis (*N*_*electrode*_ = 46 in this study). The embedded time lag was chosen based on previous studies demonstrating accurate decoding of movement kinematics and muscle activation from low-frequency EEG (28, 29). The feature vectors for each time point were labeled as class 0 (swing phase) and class 1 (stance phase). To avoid including EEG activity during different gait phases in a feature vector, we excluded the feature vectors at 10-time samples from both ends of the phases. Dataset imbalances pose difficulties to most classification analyses (73). For example, test samples belonging to the smaller class tend to be misclassified more often than those belonging to the larger class. Therefore, the number of feature vectors (time points) of the stance phase was randomly thinned out to match that of the swing phase for each stride because the duration of the stance phase was longer than that of the stance phase.

We attempted to decode the gait phase information from the time-embedded EEG data using a rLDA (“fitdiscr” function in MATLAB). rLDA shows better performance by overcoming problems such as multicollinearity and overfitting inherent in a high-dimensional dataset (the time embedded EEG dataset in this study) compared to conventional LDA (33, 74). The regularization parameter lambda was computed using a Bayesian optimization method (“OptimizeHyperparameters” option for “fitdiscr” function in MATLAB). rLDA finds an optimized weight vector *W*_*ij*_, which consists of weights at electrode *i* and time lag *j*; it is a projection vector that optimally separates the two classes (stance and swing phases). By projecting the high dimensional feature vector (the time embedded EEG data) unto a line (i.e., one-dimensional subspace), the neural decoders classify gait phases based on the cortical activity. The percentage of correctly decoded time samples (i.e., decoding accuracy) was used to quantify the decoding performance. For each subject, the decoding accuracy was calculated by 5-fold cross-validation, where the data recorded during the 6-min task were divided into 6 segments (1 min each); 5 segments were used as training data while the remaining segment was used for testing the decoding model.

To evaluate the spatial contributions of the cortical regions to the decoding of the gait phases, we calculated the contribution of each electrode from the weights of the decoding model based on previous studies (29, 35, 37):

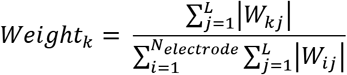

for all *k* from 1 to *N*_*electrode*_, *Weight*_*k*_ is the normalized contribution of each EEG electrode *k* against the sum of all the weights.

The EEG electrodes were assigned to 11 regions of interest (ROIs) to examine the contribution of each cortical area in the decoding model. The ROIs were left anterotemporal (FT7, FC5, T7, and C5), right anterotemporal (FT8, FC6, T8, and C6), left frontocentral (FC3, FC1, FCz, C3, C1, and Cz), right frontocentral (FC4, FC2, FCz, C4, C2, and Cz), left posterotemporal (TP7, CP5, P7, and P5), right posterotemporal (TP8, CP6, P8, and P6), left parietal (CP3, CP1, CPz, P3, P1, and Pz), right parietal (CP4, CP2, CPz, P4, P2, and Pz), left occipital (PO7, PO5, PO3, O1, and Oz), right occipital (PO8, PO6, PO4, O2, and Oz), and the central midline (FCz, Cz, CPz). The normalized weights obtained by equation 1 were averaged within each ROI.

### Statistical analysis

To test if the decoding accuracy of the neural decoders exceeded the chance level, we examined the confidence intervals using a randomization test (36). In this test, the labels of the feature vectors were randomly permuted over the time points for each subject and condition. The decoding accuracy was calculated. This procedure was iterated 500 times. The upper end of the 95 percentile was chosen as the significance level for each subject and condition.

The differences among the decoding accuracies of the neural decoders during the AO+MI, AO, and MI tasks were compared using two-tailed paired t-tests. Normality was confirmed by the Lilliefors test. The p-values from the paired t-test were corrected using the false discovery rate (FDR) for multiple comparisons (75). Additionally, the differences between the decoding accuracies of the swing and stance phases of each condition were compared using two-tailed paired t-tests. Normality was confirmed by the Lilliefors test.

The mean weights of the model of the ROIs were compared. Because normality was not observed in all the cases (tested by the Lilliefors test), the mean weights were compared using a permutation test involving 1000 random permutations (76) with FRD correction.

## Acknowledgments

This work was supported by a Grant-in-Aid for the Japan Society for the Promotion of Science Fellows (JSPS, #18J01286) to H.Y., Grant-in-Aid for Scientific Research (A) from JSPS to K.N. (#18H00818), and the Core Research for Evolutional Science and Technology (CREST) from Japan Science and Technology Agency (JST) to K.W. and K.N. (#JPMJCR14E4).

---

**Table S1.** Summary of statistical analyses for the comparisons of the decoding accuracies of the conditions. Related to Figure 4.

| Method | Pair | Value |
| --- | --- | --- |
| ANOVA | | $F_{2,24} = 8.17$, $p = 0.0020$ |
| Paired t-test with FDR correction | AO+MI vs. AO | $p = 0.011$ (0.0076 [before correction]) |
| Paired t-test with FDR correction | AO+MI vs. MI | $p = 0.0016$ (0.00054 [before correction]) |
| Paired t-test with FDR correction | AO vs. MI | $p = 0.26$ (0.26 [before correction]) |

**Table S2.** Summary of decoding accuracies and significance levels of the participant during each condition. Related to Figure 4.

| Participant ID | Experimental Condition |  |  |  |  |  |
| --- | --- | --- | --- | --- | --- | --- |
|  | AO+MI |  | AO |  | MI |  |
|  | Decoding accuracy (DA) | Significance level (SL) | DA | SL | DA | SL |
| 1 | 67.94 | 51.45 | 67.97 | 51.46 | 59.35 | 51.40 |
| 2 | 74.45 | 51.46 | 70.28 | 51.47 | 63.50 | 51.30 |
| 3 | 65.88 | 51.51 | 68.76 | 51.60 | 60.94 | 51.35 |
| 4 | 71.60 | 51.47 | 51.71 | 51.51 | 73.58 | 51.35 |
| 5 | 60.63 | 51.42 | 56.31 | 51.50 | 64.11 | 51.39 |
| 6 | 70.28 | 51.50 | 70.21 | 51.47 | 60.18 | 51.38 |
| 7 | 67.45 | 51.54 | 61.54 | 51.49 | 52.48 | 51.24 |
| 8 | 69.56 | 51.41 | 54.79 | 51.39 | 55.74 | 51.23 |
| 9 | 78.28 | 51.41 | 75.65 | 51.53 | 64.76 | 51.24 |
| 10 | 83.14 | 51.45 | 80.46 | 51.46 | 59.32 | 51.29 |
| 11 | 85.43 | 51.42 | 75.15 | 51.41 | 77.53 | 51.25 |
| 12 | 77.79 | 51.43 | 67.65 | 51.52 | 67.85 | 51.27 |
| 13 | 63.12 | 51.44 | 60.74 | 51.44 | 56.59 | 51.27 |
| Mean | 71.97 | 51.46 | 66.25 | 51.48 | 62.76 | 51.30 |
| SD | 7.21 | 0.04 | 8.35 | 0.05 | 6.76 | 0.06 |

**Table S3.**
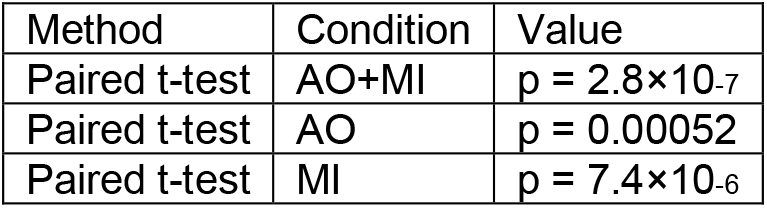
Summary of statistical analyses for the comparisons of the decoding accuracies of the stance and the swing phases of each condition. Related to Figure 5.

**Table S4.** Summary of statistical analyses for the comparisons of the contributions of the ROIs to the decoding and the mean of all electrodes during AO+MI. Related to Figure 7B.

| Method | ROI | Value (before correction) |
| --- | --- | --- |
| Permutation test with FDR correction | Left anterotemporal | p = 0.029 (0.0080) |
| Permutation test with FDR correction | Left frontocentral | p = 0.99 (0.99) |
| Permutation test with FDR correction | Right frontocentral | p = 0.65 (0.41) |
| Permutation test with FDR correction | Right anterortemporal | p = 0.36 (0.19) |
| Permutation test with FDR correction | Left posterotemporal | p = 0.99 (0.91) |
| Permutation test with FDR correction | Left parietal | p = 0.73 (0.53) |
| Permutation test with FDR correction | Right parietal | p = 0.10 (0.038) |
| Permutation test with FDR correction | Right posterotemporal | p = 0.77 (0.63) |
| Permutation test with FDR correction | Left occipital | p = 0.029 (0.0060) |
| Permutation test with FDR correction | Right occipital | p = 0.029 (0.0070) |
| Permutation test with FDR correction | Central midline | p = 0.18 (0.084) |

**Table S5.** Summary of statistical analyses for the comparisons of the contributions of the ROIs to the decoding and the mean of all electrodes during AO. Related to Figure 7C.

| Method | ROI | Value (before correction) |
| --- | --- | --- |
| Permutation test with FDR correction | Left anterotemporal | p = 0.0066 (0.0030) |
| Permutation test with FDR correction | Left frontocentral | p = 0.014 (0.0090) |
| Permutation test with FDR correction | Right frontocentral | p = 0.0037 ($1.0 \times 10^{-4}$) |
| Permutation test with FDR correction | Right anterortemporal | p = 0.67 (0.67) |
| Permutation test with FDR correction | Left posterotemporal | p = 0.13 (0.098) |
| Permutation test with FDR correction | Left parietal | p = 0.0037 ($1.0 \times 10^{-4}$) |
| Permutation test with FDR correction | Right parietal | p = 0.26 (0.24) |
| Permutation test with FDR correction | Right posterotemporal | p = 0.13 (0.11) |
| Permutation test with FDR correction | Left occipital | p = 0.0037 ($1.0 \times 10^{-4}$) |
| Permutation test with FDR correction | Right occipital | p = 0.0073 (0.0040) |
| Permutation test with FDR correction | Central midline | p = 0.0037 ($1.0 \times 10^{-4}$) |

**Table S6.** Summary of statistical analyses for the comparisons of the contributions of the ROIs to the decoding and the mean of all electrodes during MI. Related to Figure 7D.

| Method | ROI | Value (before correction) |
| --- | --- | --- |
| Permutation test with FDR correction | Left anterotemporal | p = 0.010 (1.0×10<sup>-4</sup>) |
| Permutation test with FDR correction | Left frontocentral | p = 0.95 (0.71) |
| Permutation test with FDR correction | Right frontocentral | p = 0.96 (0.96) |
| Permutation test with FDR correction | Right anterortemporal | p = 0.15 (0.056) |
| Permutation test with FDR correction | Left posterotemporal | p = 0.67 (0.30) |
| Permutation test with FDR correction | Left parietal | p = 0.96 (0.90) |
| Permutation test with FDR correction | Right parietal | p = 0.95 (0.78) |
| Permutation test with FDR correction | Right posterotemporal | p = 0.95 (0.76) |
| Permutation test with FDR correction | Left occipital | p = 0.022 (0.0040) |
| Permutation test with FDR correction | Right occipital | p = 0.033 (0.0090) |
| Permutation test with FDR correction | Central midline | p = 0.96 (0.53) |

## Notes

### Competing Interest Statement

The authors have declared no competing interest.

